# Decoding ultrasensitive self-assembly of the calcium-regulated *Tetrahymena* cytoskeletal protein Tcb2 using optical actuation

**DOI:** 10.1101/2025.05.26.656216

**Authors:** Nithesh P. Chandrasekharan, Xiangting Lei, Jerry Honts, Saad Bhamla, Scott M. Coyle

## Abstract

EF-hand calcium binding proteins are key macromolecular components of many unique filament systems and ultrafast contractile structures found in protists. However, our biochemical understanding of these cytoskeletal systems has been hindered by the need for assays that can controllably generate spatiotemporal calcium dynamics to probe their behavior. Here, we define the quantitative requirements for calcium-dependent self-assembly of the *Tetrahymena* cortical cytoskeletal protein Tcb2 using a microscopy-based spatiotemporally controlled optical calcium release assay. Light-driven uncaging of the photolabile calcium chelator DMNP-EDTA stimulates rapid localized self-assembly of Tcb2 into micron-scale protein networks. We quantify how the growth, size, and lifetime of Tcb2 networks is controlled by the duration and intensity of the applied light pulse. Incorporating the fluorescent calcium indicator Rhod-5N allows inference of the spatiotemporal distribution of calcium-bound Tcb2 monomers during the reaction and identifies a sharp, ultrasensitive transition to Tcb2 self-assembly. By applying this assay to mutants in Tcb2’s four EF hand domains, we show that D184 is the key calcium binding site that licenses Tcb2 for self-assembly and define quantitative roles for other binding sites in tuning Tcb2’s calcium-responsiveness. Our approach reveals a rich space of structures and regulation available to a single-protein system through coupling calcium-binding to ultrasensitive self-assembly, opening new paths forward to understanding other protist filament networks and contractile myonemes.

## Introduction

Calcium ions are ubiquitous players in the regulation of cell structure and mechanics^1–4^. By sequestering calcium in stores and pumping it out of the cytoplasm, cells position themselves to regulate cell-biological processes through rapid influx of these ions in response to internal or external cues^4–8^. For example, muscle cell contraction is actuated when calcium ions released from the sarcoplasmic reticulum bind tropomyosin, triggering conformational changes that expose binding sites for myosin motors on actin filaments^9–11^. Likewise, the binding of calcium ions to the signaling protein calmodulin regulates the activity of Myosin Light Chain Kinase which controls contractility in smooth muscle and other non-muscle metazoan cell types^12–14^ . In each of these cases, calcium acts as a signaling regulator of proteins whose downstream activity influences the behavior of ATP-powered contractile systems.

Calcium ions can also play a more direct role in the assembly and contraction of cellular structures^15–19^. Indeed, many single-celled protists contain exotic filament systems that underlie sophisticated cell structures and morphologically defined behaviors that are assembled and controlled in part through the action of calcium-binding proteins^20–24^. For example, the peritrich ciliate Vorticella^25,26^ and the heterotrich ciliate Spriostumum^23,25^ both contain contractile structures called myonemes that facilitate ultrafast contractility (instantaneous speeds on the order of mm/s) that are thought to enable relocation of feeding currents and predatory escape respectively. In these systems, centrin-family or spasmin-family EF-hand calcium binding proteins assemble on scaffold proteins and directly trigger contractility upon calcium binding following influx from internal and external stores^22,25–27^. Reversibility of the contracted structure is typically slow and depends indirectly on ATP through the action of calcium pumps, which work to shift the equilibrium state of the structure back to a relaxed confirmation and reset the system^25,27^. While there is increasing interest in the biochemical and biophysical mechanisms driving ultrafast myoneme contraction^25,28–30^, a detailed understanding has been hindered by challenges in reconstituting actuatable version of these systems, given the size and complexity of the multi-component protein structures involved and the need for dynamic spatiotemporal control over the calcium ions that actuate and regulate their activity.

Within the ciliate *Tetrahymena thermophila*, Tcb2 (also known as TCBP-25) forms a major component of the calcium-sensitive cortical cytoskeleton (Fig. 1A)^31–33^. Like myoneme-associated centrins and spasmins, Tcb2 contains a domain architecture consisting of two calcium binding domains, each of which can bind two calcium ions through a pair of EF-hand motifs^34–36^. However, Tcb2 is not known to associate with canonical centrin-binding scaffold proteins from the Sfi1 family^37^. Instead, concentrated solutions of purified recombinant Tcb2 alone will rapidly self-assemble into higher order protein networks, contractile gels, or aggregates in the presence of calcium that are visible by light microscopy^35,38^. These features make Tcb2 a potential single-component model system for biochemical and biophysical investigations into the calcium-regulated assembly of micron-scale protein structures and networks common to protozoan cytoskeletons. However, the protein’s rapid assembly makes its dynamic behavior difficult to image and study quantitatively, as introduction of calcium through manual pipetting is difficult to control, creating mixing artifacts and gradients that muddle observation and interpretation^34–36^.

**Figure 1.**
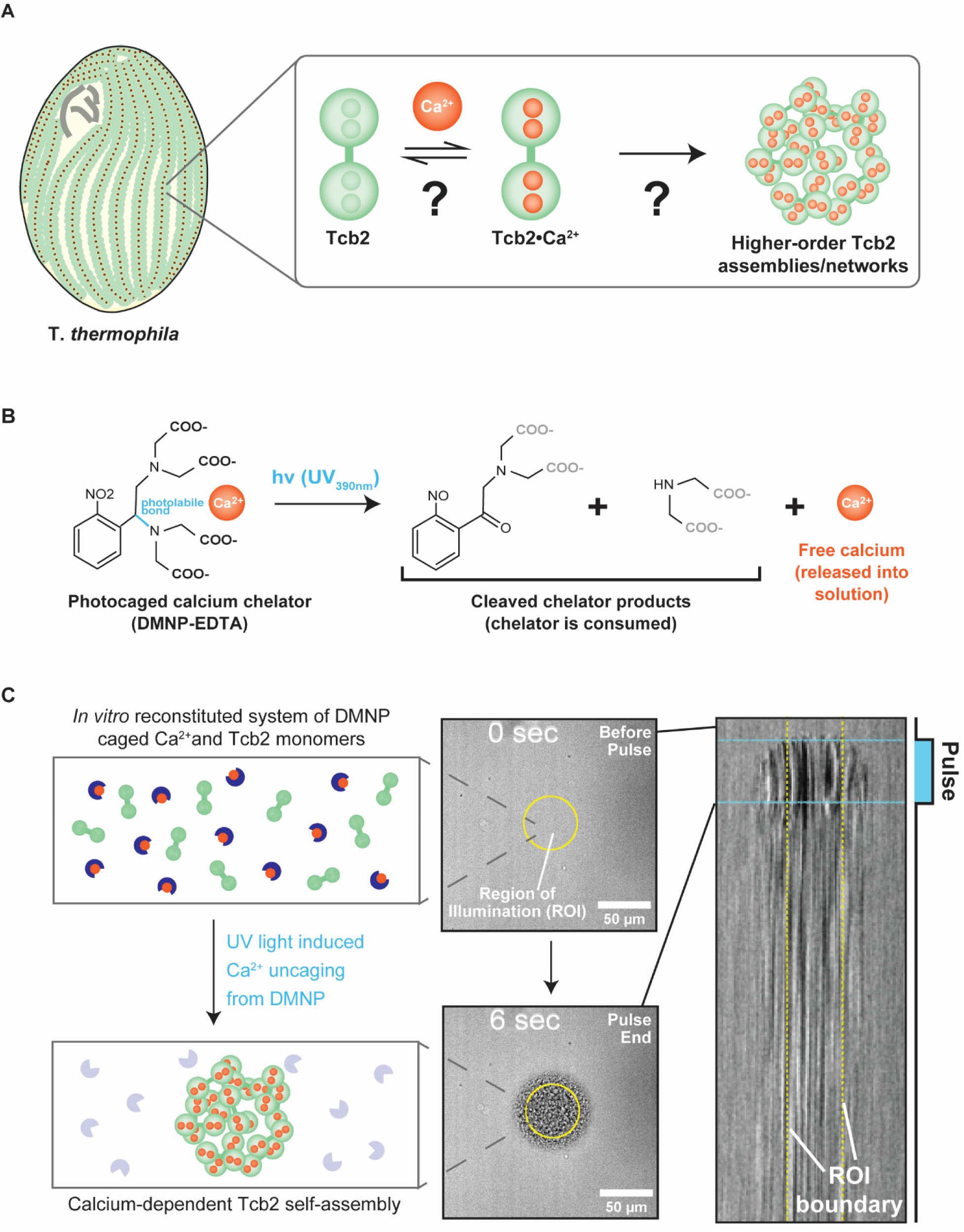
An optical actuation system for quantitative characterization of calcium-dependent Tcb2 self-assembly. A) Tcb2 part of cortical lattice layer surrounding basal bodies of cytoskeleton. Tetrahymena calcium binding protein 2 (Tcb2) forms self-assembled networks induced by calcium ions. B) Design for Photo-stimulation *In vitro* Calcium Hyper-fast Uncaging (P*I*CHU) assay. Caged calcium chelator, DMNP-EDTA, contains a photolabile bond that breaks upon UV390nm stimulation to release its caged calcium ions. Uncaged calcium ions bind to EF-hands of Tcb2 with varying affinity. With our optical setup, we can trigger uncaging and thus calcium induced Tcb2 self-assembly fiber formation visualized using brightfield imaging. C) Brightfield kymograph depicting growing Tcb2 self-assembly over time. Yellow dotted lines indicating ROI boundary.

Here, we expand on our recently developed optically controllable Tcb2 actuation assay^38^ to quantitatively probe the biochemical mechanisms and constraints on calcium-dependent Tcb2 self-assembly. We use the UV-sensitive photocaged calcium chelator DMNP-EDTA^39^ to controllably project micron-scale calcium inputs onto Tcb2, allowing us to directly relate different patterns of spatiotemporal calcium dynamics to the formation and behavior of higher-order Tcb2 network assemblies. By combining this approach with the fluorescent calcium-indicator Rhod-5N^40^ , we could quantitatively disentangle calcium-loading of soluble Tcb2 monomers in solution from assembly into higher-order networks. Using this approach, we identify a threshold concentration of calcium loading needed for actuation that defines a sharp, ultrasensitive boundary between soluble and assembled Tcb2 states. Site-directed mutagenesis of different calcium binding sites within Tcb2’s EF hand domains identified a single calcium binding site (EF4: D184) as absolutely critical for self-assembly, while other binding sites appear to tune the critical concentration and dynamics of assembly. Together, we clarify how the coupling of calcium binding to ultrasensitive self-assembly can regulate the size and dynamics of Tcb2 structures. More generally, our optical assay provides a platform for actuating and probing the biochemical and biophysical mechanisms of calcium-responsive myonemes and filaments with the spatiotemporal control needed to fully understand their remarkable functions and capabilities.

## Results

### An optical actuation system for quantitative characterization of calcium-dependent Tcb2 self-assembly

Tcb2 is a calcium-binding protein that forms a key element of the *Tetrahymena* cortical cytoskeleton^37^ . In the presence of calcium, concentrated solutions of recombinant Tcb2 form fibrous protein networks that are visible by brightfield microscopy^35,38^ . A proposed model for Tcb2 *in vitro* network formation is that calcium-bound Tcb2 monomers self-associate to form higher order assemblies (Fig. 1A)^34,35,38^. However, little is known about the specific quantitative calcium requirements for Tcb2 network formation nor how each calcium binding site in the Tcb2 protein contributes to self-assembly. This is in part due to the challenges of triggering Tcb2 network formation in a controlled manner, as introduction of calcium ions through pipetting leads to rapid non-homogenous network formation driven by mixing artifacts^35,38^.

To develop an assay that provides spatial and temporal control over calcium-dependent Tcb2 network formation, we took advantage of the photocaged calcium chelator DMNP-EDTA^39^ . The K_D_ of DMNP-EDTA for Ca^2+^ increases from 5nM to 3mM upon photolysis at UV_390nm_, allowing for optically controlled release of calcium (Fig. 1B). We hypothesized that the role of calcium dynamics on Tcb2 network formation could be quantitatively investigated microscopically using a digital mirror device (DMD) to locally and quantitatively uncage DMNP-EDTA in Tcb2-containing solutions. We previously applied a similar method to investigate the mechanical and material properties of Tcb2 networks *in vitro* under saturating conditions^38^.

To test, we prepared a reaction solution consisting of 500µM caged Ca·DMNP-EDTA, 250µM Tcb2, and 100µM EGTA chelator and mounted 1μl in a fixed height chamber for imaging and stimulation with a 40X objective. These concentrations approach the high Tcb2 levels observed physiologically in cells^37^, while the inclusion of EGTA chelator stabilizes Tcb2 protein in solution and buffers actuation of the system. These reaction solutions showed no self-assembly in the absence of light. However, upon DMD stimulation of a region of interest (ROI) within the sample using UV_390nm_ light (pulse parameters: 50µm ROI diameter, 5sec pulse, 100% intensity), we observed formation of Tcb2 protein networks within milliseconds (Fig. 1B, Movie S1). During light stimulation, Tcb2 networks grow outwards and begin to expand beyond the radius of the ROI, implying that the diffusion of released calcium plays a role in the growth of Tcb2 networks in the *in vitro* assay (Fig. 1C). Once the pulse ended, Tcb2 network growth stopped and the visible boundary of the network partially receded at its edges, leaving a final Tcb2 network structure that was stable and persisted for the duration of our imaging experiment. These data confirm that Tcb2 networks can be triggered and visualized *in vitro* by optically driven DMNP-EDTA calcium release.

### Tcb2 self-assembly is sensitive to different optical calcium-release protocols

A strength of our optical strategy for triggering Tcb2 self-assembly is that the levels and rates of calcium release during the reaction can in principle be controlled by varying the light intensity (0-100%) and duration of the applied UV light within the ROI used for stimulation. This provides a means of asking how Tcb2 network formation responds to different kinds of spatiotemporal calcium dynamics (Movie S2). To quantify how different pulse protocols affect Tcb2 network formation, we developed an image analysis pipeline that detects and tracks the radial growth of the visible Tcb2 network boundary over time during a stimulation experiment (Fig. 2A).

**Figure 2:**
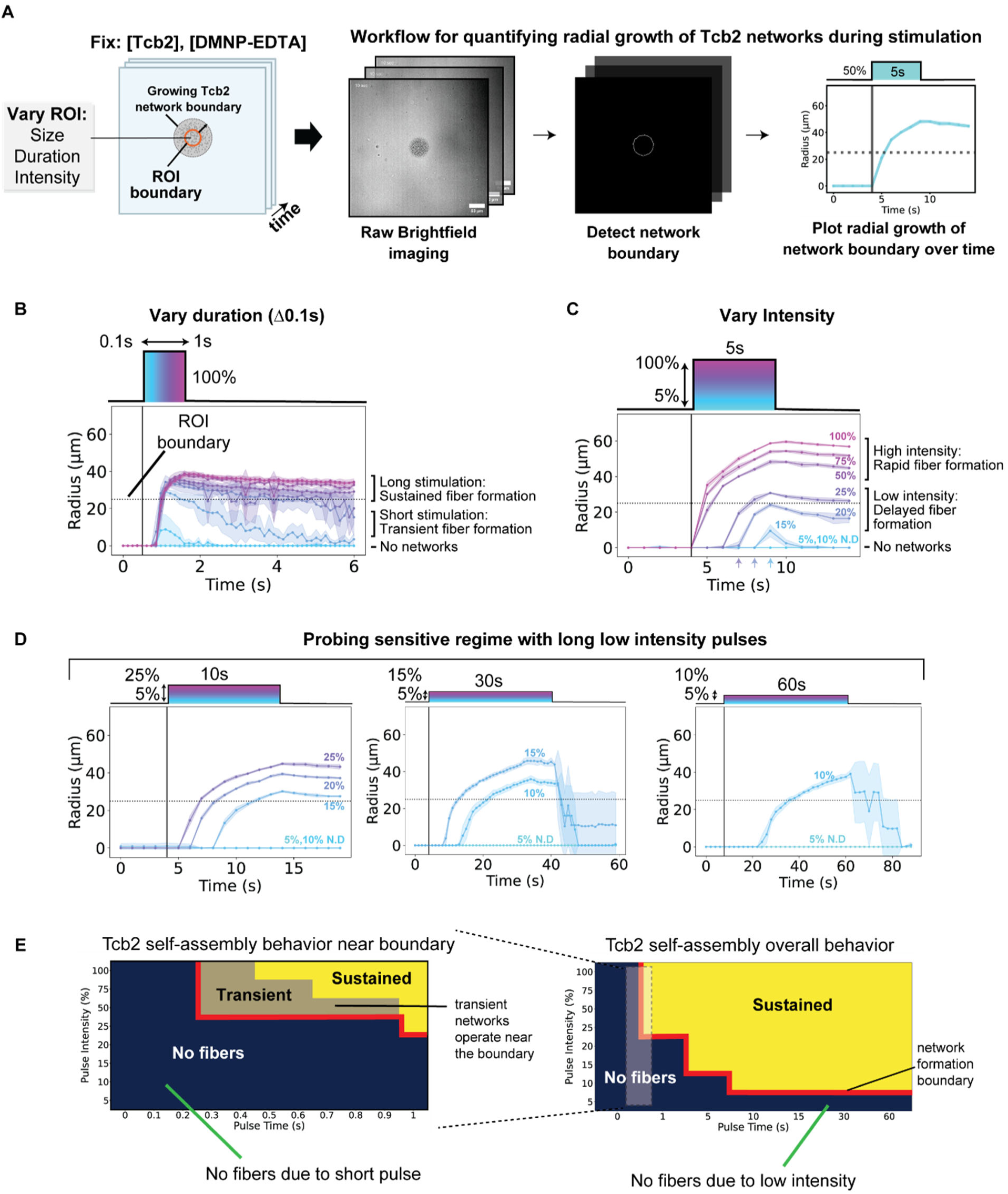
Tcb2 self-assembly is sensitive to different optical calcium-release protocols. A) Workflow for quantifying radial growth of Tcb2 self-assembly fiber formation. Contours of Tcb2 fiber growth are obtained using our FiJI/Python pipeline and the radius of each contour is plotted over time. B) Varying pulse parameters show different Tcb2 fiber formation phenotypes. Short high intensity pulses result in transient fibers while longer pulses lead to sustained fibers. Lowering the intensity leads to a delay in fiber formation. C) Probing the sensitive region of Tcb2 fiber formation by using long low intensity pulses to determine when fiber formation is triggered. D) Portfolio showing sensitivity of Tcb2 fiber formation based on the varying classes of calcium releases due to tunability of pulse parameters.

Using this approach, we first fixed illumination intensity at 100% and asked how the duration of a pulse affected Tcb2 network formation. No network formation was detected for short pulses <300ms (100ms, 200ms). In contrast, pulses longer than 300ms began to show detectable network formation, with all pulses growing at an initial rate of 86μm/s^-1^ ± 10μm/s^-1^ (n=35). Short pulses (300ms, 400ms) led to networks that were transient in character, beginning to disassemble significantly following the end of the pulse. However, for pulses >500ms, networks persisted and were sustained well after the pulse was completed, showing little to no disassembly. Tcb2 network formation is thus sensitive to the duration of the calcium release at a fixed intensity.

We next fixed the duration of our light pulse to 5s and asked what the effects of pulse intensity were on Tcb2 network formation. Pulses >50% generated Tcb2 networks almost immediately (within 200ms), with the final radius of the Tcb2 network scaling with the intensity of the applied pulse (Fig. 2B). Interestingly, pulses <50% also formed networks, but showed a significant delay between initiation of the pulse and appearance of the network. For example, a 25% intensity pulse took 2.4s± 0.5s (n=8) to produce a detectable network, 20% intensity pulse took 3.4s± 0.5s (n=8), and pulses <15% did not produce networks during the pulse. Thus, the delay between onset of stimulation and appearance of network appeared to correlate with the pulse intensity. To explore this further, we tested whether 5% and 10% pulses could form networks if longer stimulation times we used. We found that 10% intensity pulses could indeed produce Tcb2 networks after 13±3.8 s (n=8), while we were unable to detect network formation for 5% pulses even after 60s of stimulation (Fig. 2C).

The key observations from this panel of stimulation experiments can be summarized in a phase portrait that maps the optical actuation protocol used to the Tcb2 network formation state observed (sustained, transient, none) (Fig. 2D). This phase portrait reveals a clear and intuitive boundary in our experimental calcium uncaging strategy separating pulses that can drive self-assembly and those that cannot. Short pulses require high light intensities to drive Tcb2 network formation, while longer pulses at lower intensity can also support Tcb2 assembly, but with considerable delays. Transient network formation was associated with pulse protocols operating near the network formation boundary. This suggests that if we can quantify the spatiotemporal calcium dynamics generated by these different pulses, we can likely determine the calcium requirements for Tcb2 self-assembly and the phenomena we observe.

### Quantifying spatiotemporal calcium uncaging dynamics using Rhod-5N fluorescence

We next sought to understand how the different light stimulation protocols we used to actuate Tcb2 self-assembly affect the underlying spatiotemporal dynamics of calcium release that triggers this process. To this end, we used Rhod-5N^40^, a low affinity fluorometric BAPTA-based calcium indicator, to visualize calcium uncaging dynamics in our assay in real time (Fig. 3A). We selected Rhod-5N due to 1) its red-shifted excitation and emission wavelengths (Ex/Em of Ca^2+^–bound form: 551/576nm) providing good separation from the UV_390nm_-activated DMNP-EDTA caged chelator and 2) its high K_D_ for calcium (∼320μM) allowing it to effectively report on the calcium concentrations needed to load highly concentrated solutions of Tcb2 (250μM).

**Figure 3.**
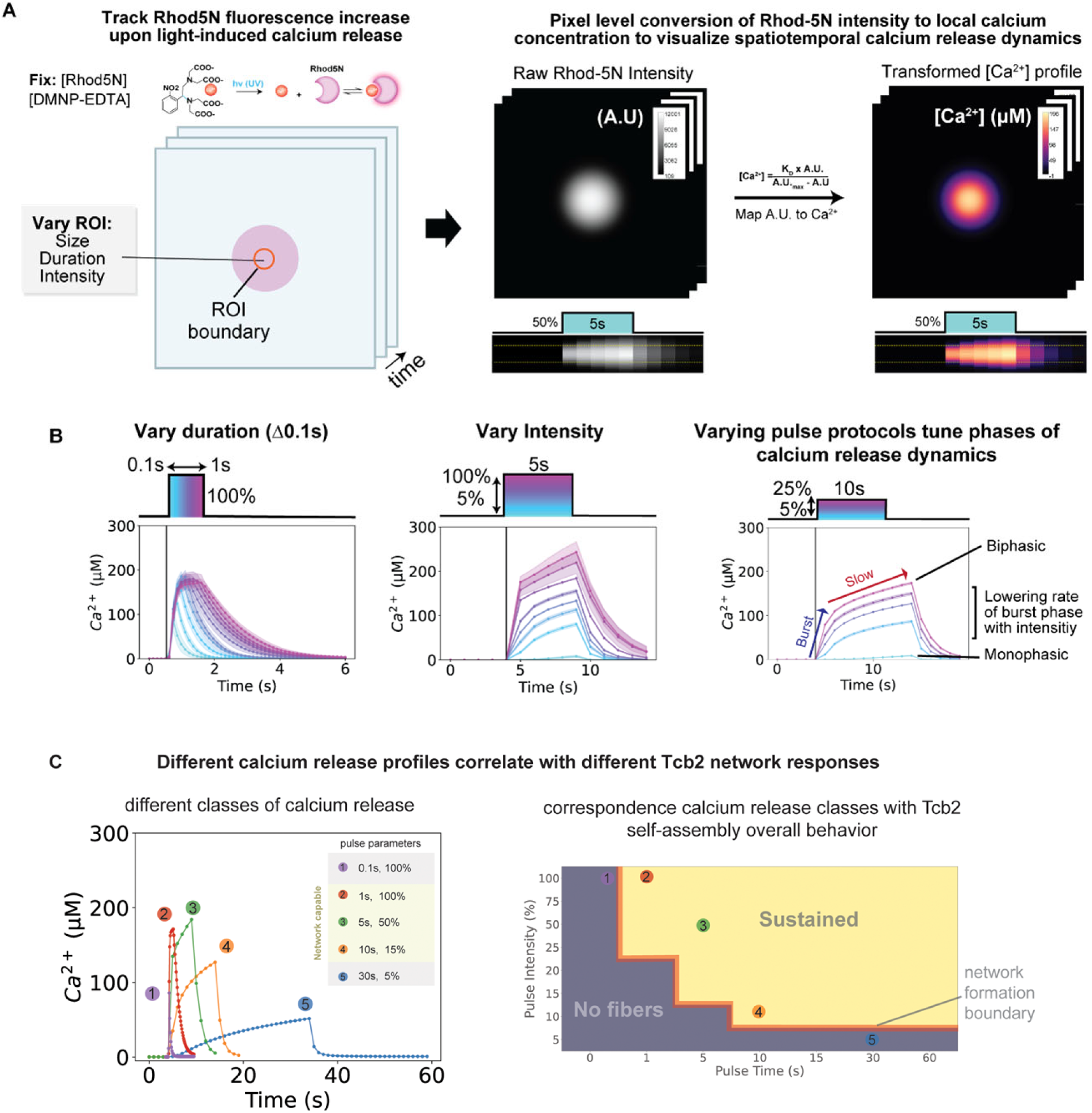
Quantifying spatiotemporal calcium uncaging dynamics using Rhod-5N fluorescence. A) Workflow for imaging and quantification of Rhod-5N fluorescence during stimulation. Calibration equation generated from Rhod-5N standard curve (SI Fig) used to convert intensity to [Ca^2+^]_free_ at pixel level. Kymographs show the spreading of intensity (indicating calcium binding to Rhod-5N) over time during stimulation and decay after pulse end. B) Varying pulse parameters describes calcium release profiles. Short high intensity pulses create transient bursts of calcium release. Increasing pulse time leads to a transient burst then followed by slow steady release. Lowering intensity reduces the burst phase and ultimately only shows a slow steady release. C) The 3 classes of calcium release can be related to the different Tcb2 fiber assembly profiles. Short high intensity pulse with a quick release of calcium result in transient fibers. Increasing the pulse time increases the amount of calcium released and thus sustained fibers. Lowering the intensity releases less calcium and not sufficient for fiber formation. However, with increasing pulse time, enough calcium is eventually released to result in delayed fibers.

Using the same buffer system as in our previous assays but substituting 25µM Rhod-5N for Tcb2, we first tested whether light-driven DMNP-EDTA uncaging produced a spatiotemporal change in the intensity of the Rhod-5N dye as judged by fluorescence microscopy. In the absence of light, Rhod-5N signal was dim and uniform throughout the field of view. However, upon a localized stimulation of DMNP-EDTA within a 50μm ROI (pulse parameters: 50%, 5s), we observed an immediate increase in the Rhod-5N intensity (Fig. 3A). During light stimulation, the intensity increased over time and spread radially outwards and expanded beyond the ROI via diffusion. After the pulse, Rhod-5N intensity stopped increasing but continued to spread spatially outwards from the ROI by diffusion. Simultaneously, the fluorescence intensity within the field of view decayed back to baseline levels, owing to competition between Rhod-5N and the higher affinity calcium chelator EGTA present in the assay buffer. These data indicate that our optical uncaging strategy produces a spatiotemporal calcium signal that can be visualized everywhere within the field of view using Rhod-5N fluorescence.

To convert the observed pixel-level Rhod-5N intensities into estimates of local [Ca^2+^], we collected full-field images of calcium standards (final [Ca^2+^] from 2µM to 25mM) in the presence of 25µM Rhod-5N in our assay buffer using the same imaging chambers and optical configuration we used for our Tcb2 actuation experiments. The resulting intensity profiles were fit to a binding curve to generate a calibration equation that identifies the linear range of the Rhod-5N under our assay conditions (Fig S1). Rearranging this equation allows us to convert pixel-level Rhod-5N fluorescent intensities into estimates of the local [Ca^2+^] all throughout the field of view for different stimulation profiles (Fig. 3A).

Using this strategy, we can visualize and quantify the spatiotemporal calcium release profile associated with any optical stimulation protocol used in our previous Tcb2 experiments from Figure 2. While these experiments provide spatiotemporal information about the local calcium concentrations everywhere within the field of view, to gain intuition about the behavior of different pulses we specifically plotted the maximum calcium concentration over time observed in the center of the ROI to compare across all pulse protocols. As in our earlier Tcb2 assays, we first fixed an illumination intensity at 100% and asked how the duration of a pulse affected calcium release dynamics (0.1-1s, Δ0.1s). All pulses, even the shortest 100ms pulse, produced immediate and readily detectable calcium release (Fig. 3B). During early timepoints of stimulation (<500ms), each of the pulses released calcium at a similar rate (350μM/s^-1^ ± 35μM/s^-1^, n=50). However, at longer timepoints (>500ms) this initial rapid release of calcium slowed significantly (19μM/s^-1^ ± 9μM/s^-1^, n=25). After completion of the pulse, calcium levels decayed back to baseline at timescales that appeared to vary directly with the duration of stimulation.

We next fixed the duration of our pulse to 5s and asked what the effects of pulse intensity (5-100%) were on calcium release dynamics (Fig. 3B). For high intensity pulses (>50%), we observed a rapid burst of calcium release over the first second followed by a slower sustained release rate for the duration of the pulse. This is similar to what was observed for 100% intensity pulses >500ms. However, as the pulse intensity decreased further, the initial rates of calcium release became slower, and the burst behavior became less pronounced (Fig 3B, Fig S2). Indeed, at intensities <15% the calcium release rate became monophasic and was mostly linear throughout the duration of the experiment.

Together, our empirical observations suggest that spatiotemporal effects of the localized light-driven DMNP-EDTA uncaging reaction and diffusion together impact the emergent calcium dynamics we generate in our assay. For high intensity pulses, a biphasic calcium release behavior is observed in which an initial burst of calcium is released as DMNP-EDTA is rapidly consumed within the ROI, followed by slower calcium release as rate-limiting step shifts from photolysis to diffusion of caged DMNP-EDTA molecules into the ROI from the surroundings. For lower intensity pulses, the rate of DMNP-EDTA uncaging is slow enough that it does not significantly deplete the available supply faster than can be replenished by diffusion, allowing for a more consistent rate of release throughout the experiment (Fig. 3B).

A key takeaway from these observations is that our optical uncaging assay generates different classes of calcium release dynamics probe Tcb2 network formation in different ways (Fig. 3C). Brief, rapid bursts of calcium support the rapid assembly of short-lived transient Tcb2 networks we observed on our phase diagram (see Fig. 2D). Similarly, sustained release of high calcium supported rapid formation of larger, more sustained Tcb2 networks. In contrast, slow calcium release rates with low calcium levels led to delays in the onset of Tcb2 network formation or failed to support self-assembly at all. Together, our calibration of calcium release using Rhod-5N clarifies the calcium dynamics at play in our optical assay and provides a portfolio of spatiotemporal dynamics to investigate the calcium requirements for Tcb2 self-assembly.

### A combined Tcb2 / Rhod-5N actuation assay reveals a sharp, threshold level of calcium required for self-assembly

Having seen that optically-triggered Tcb2 network formation is sensitive to the intensity and duration of the applied pulse of light, and that different pulse protocols produced marked different spatiotemporal calcium release profiles as judged by the calcium indicator dye Rhod-5N, we next aimed to combine these approaches together into a single assay to directly explore the relationship between calcium binding to Tcb2 and self-assembly. To this end, we exploited competition between Tcb2 and Rhod-5N for binding to calcium to estimate the spatiotemporal distribution of calcium bound to Tcb2 (henceforth: Ca·Tcb2) during network formation.

To explore this, we prepared Rhod-5N reaction solutions in the same buffer systems as previously with or without 250μM Tcb2. As before, Rhod-5N fluorescence was dim in the absence of light simulation, and Tcb2 containing solutions were stable in the absence of UV light. We then stimulated each solution with a test light pulse (50µm, 5s, 50% intensity) and followed the spatiotemporal distribution of Rhod-5N fluorescence in the field of view over time. As before, in the absence of Tcb2, Rhod-5N fluorescence rapidly increased during the pulse and spread via diffusion and decayed back to baseline after light simulation ended. In the presence of Tcb2, Rhod-5N fluorescence also increased and decreased in a qualitatively similar manner, but the observed fluorescence intensity was substantially reduced and Tcb2 networks formed simultaneously during the reaction. Thus, Rhod-5N and Tcb2 can be included in the same reaction solution and still support light-driven network assembly, and the presence of Tcb2 in the reaction reduces the apparent Rhod-5N signal through competition for the released calcium.

We took advantage of this phenomena to develop an image analysis pipeline that estimates the spatiotemporal distribution of Ca·Tcb2 at every position in the field of view. For this, we imaged Rhod-5N fluorescence during optical uncaging in the presence or absence of Tcb2 and performed a pixel-level subtraction of the image time series to measure the local change in fluorescence ΔI=I_ref_-I_exp_. Using the Rhod-5N calibration curve we established earlier, ΔI provide a first-order estimate for the local concentration of calcium loaded into Tcb2^41^. In parallel, we tracked Tcb2 self-assembly by following the visible boundary of the network using brightfield microscopy as before. This analysis pipeline allows us to track the spatiotemporal dynamics of Tcb2 calcium loading and network assembly independently during optical actuation (Movie S3).

We then used our combined assay to probe and identify the calcium loading requirements necessary to support Tcb2 self-assembly. To this end, we optically actuated Tcb2 network formation in the presence of Rhod-5N using the same suite of light stimulation pulses in Figures 2 and 3. We applied our image analysis pipeline to extract and track three metrics of interest over time: the maximum concentration Ca·Tcb2; the radius of the growing Tcb2 network boundary; and the concentration of Ca·Tcb2 at that boundary, where Tcb2 is undergoing a transition from soluble to self-assembled. These metrics over time are plotted for all pulses in Figure 4B.

**Figure 4.**
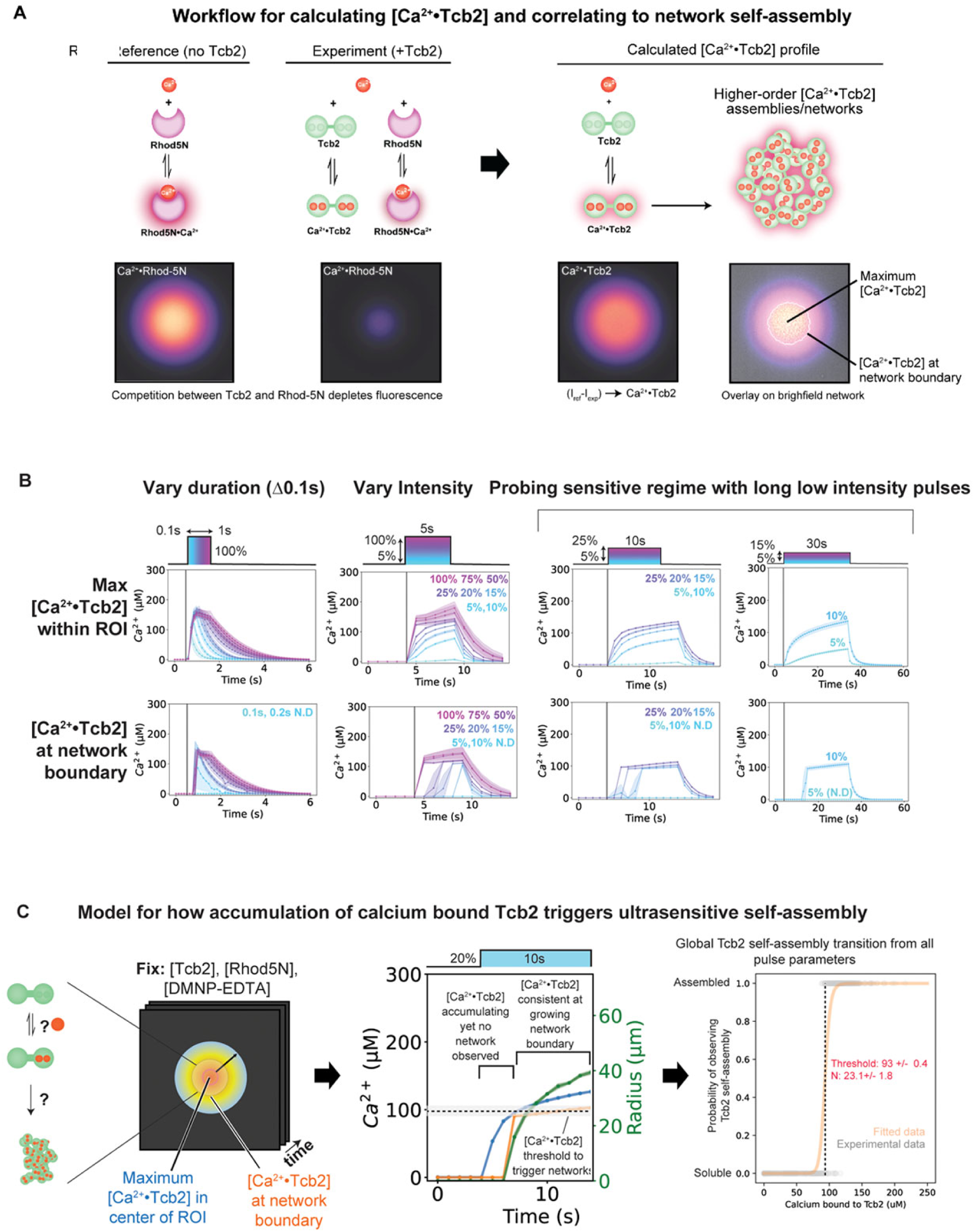
A combined Tcb2 / Rhod-5N actuation assay reveals a sharp, threshold level of calcium required for self-assembly. A) Workflow for calculating [Ca^2+^·Tcb2]_bound_ and correlating to Tcb2 self-assembly network. Rhod-5N fluorescence decreases in the presence of Tcb2. Subtracting the [Rhod5N/Tcb2·Ca^2+^] profiles from [Ca^2+^·Rhod5N] profiles provides a calculated [Ca^2+^·Tcb2]_bound_ estimate profile. To correlate to Tcb2 fiber formation, we look at two aspects, [Ca^2+^·Tcb2]_bound_ at the center of the ROI to look at the threshold that triggers fiber assembly, [Ca^2+^·Tcb2]_bound_ at the network boundary as the transition point from monomer to oligomer. B) Variation of pulse parameters (pulse time and intensity) dictate levels of calcium binding to Tcb2 and a [Ca^2+^·Tcb2]_bound_ threshold level of 114.19μM±14.78μM at the network boundary. C) Model for accumulation of [Ca^2+^·Tcb2]_bound_ at the center of the ROI up to a certain threshold concentration value which once crossed triggers fiber formation and is consistent at the network boundary when transitioning from monomer to fiber in real time.

From these data we made several key observations about the relationship between Tcb2 self-assembly and calcium loading. First, we were able to detect and track calcium loading into Tcb2 for all pulses protocols tested, including those that did not lead to observable Tcb2 network formation. This implies that Tcb2 requires a sufficiently high concentration of Tcb2 to be loaded with calcium to undergo self-assembly. To quantitatively define this requirement, we carefully examined the behavior of long duration, low-intensity pulse protocols that resulted in delayed Tcb2 network formation. For these pulse protocols, the maximum Ca·Tcb2 slowly increased at a rate set by the light intensity until network formation suddenly became detectable, at which point we consistently observed [Ca·Tcb2] to be 105μM ± 12μM (n=45) at the boundary of the Tcb2 network. To further quantify and characterize this ultrasensitive transition, we aggregated all calcium-bound Tcb2 concentration observations and their associated Tcb2 self-assembly status from 72 different experimental pulses. Fitting self-assembly as a function of [Ca·Tcb2] to a hill equation recovers a sharp transition (nH = 23.1 ± 1.8) occurring at [Ca·Tcb2] 93 ± 0.4μM (Fig. 4C).

Taken together, our data suggest a model for Tcb2 behavior in which calcium loaded Tcb2 monomers undergo a sharp, ultrasensitive transition into a self-assembled state when the system reaches a critical concentration of Ca·Tcb2 (Fig. 4C). Different pulse protocols generate different spatiotemporal calcium release profiles, which control the timescales for reaching this critical concentration in space. Pulses that release calcium at high rates cross this threshold rapidly, allowing Tcb2 networks to grow radially beyond the ROI. When the rate of release is slow, the time needed to cross this critical concentration is delayed or never reached, limiting when and where self-assembly occurs. Thus, by coupling calcium binding to a sharp self-assembly response, Tcb2 converts different spatiotemporal calcium dynamics into differences in the growth, size, and lifetime of the networks it forms.

### Distinct roles for N-terminal and C-terminal EF-hand domains in Tcb2 network formation

Tcb2 is proposed to bind calcium using distinct N-terminal and C-terminal EF-hand domains, each of which contains two putative binding sites for calcium ions^34–36^. However, the specific contributions of these different calcium binding sites to Tcb2 self-assembly are not understood. To gain insight, we predicted the structure of Tcb2 using AlphaFold and used an EF-hand consensus sequence to map the locations of conserved aspartate residues predicted to be critical for binding onto the structure (Fig 5A)^42,43^. This revealed an overall architecture for Tcb2 similar to the canonical calcium-binding protein calmodulin^44^, in which the two EF hand domains are connected by a flexible linker. In addition, Tcb2 contains an N-terminal extension predicted by AlphaFold to be disordered, which is a common feature of other centrin-family proteins associated with protist cytoskeletons^15,16^.

**Figure 5.**
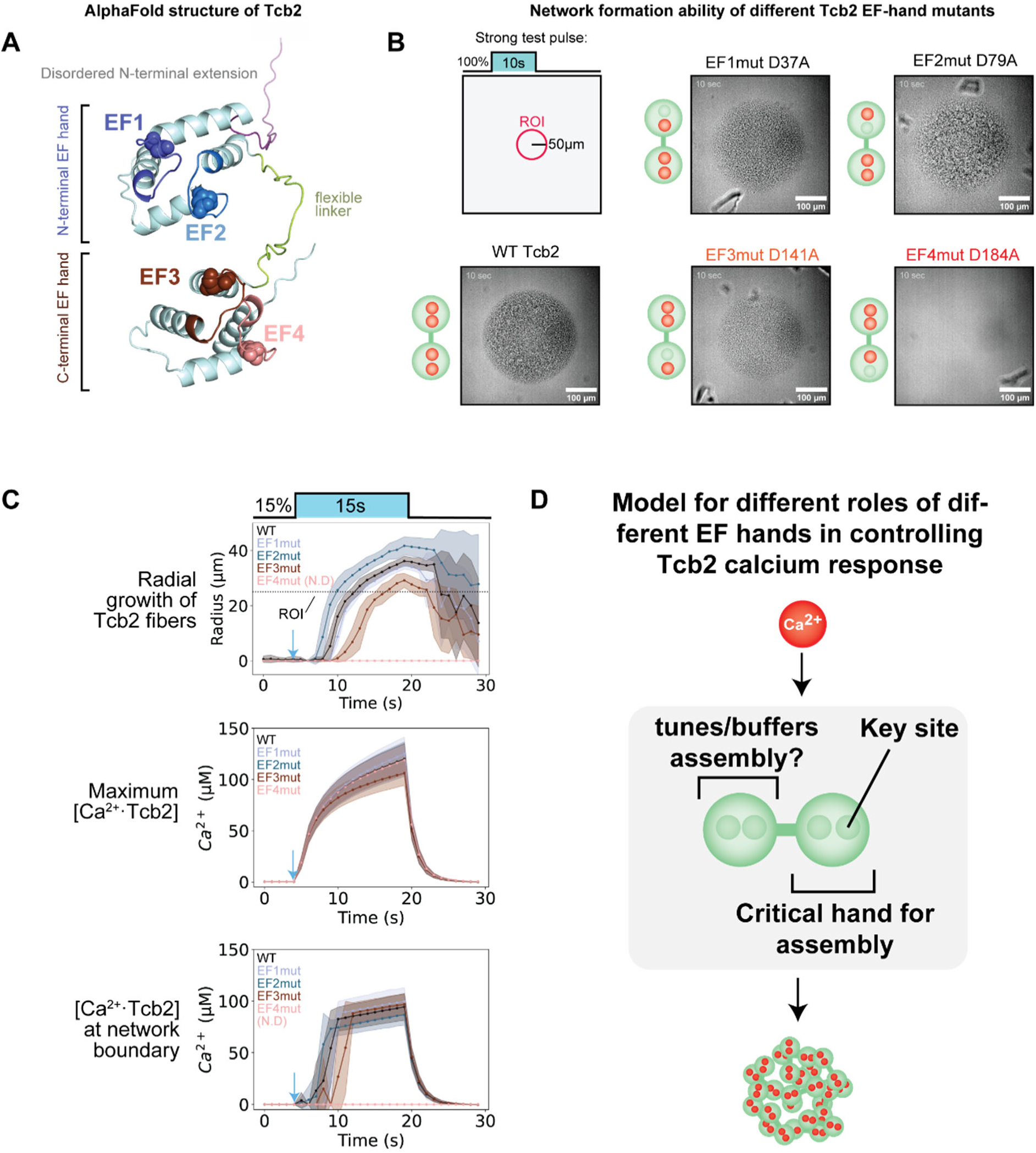
Distinct roles for N-terminal and C-terminal EF-hand domains in Tcb2 network formation. A) Predicted Tcb2 AlphaFold structure. Conserved aspartate residue for each EF-hand depicted as spheres. B) Initial phenotypes of EF-hand knockouts show detrimental effects of C-terminal EF-hand domains using a strong test pulse (10sec, 100%). EF3 KO (D141A) appeared qualitatively compromised while EF4 KO (D184A) is completely unable to form fibers. C) Using our [Ca^2+^·Tcb2]_bound_ workflow and looking at the sensitive regime teased out differences between the KOs. All KOs, including dead EF4 KO, was able to bind to Ca^2+^ like WT regardless of self-assembly. EF2 KO formed fibers faster while EF3 KO was further delayed compared to WT. At the network boundary, EF2 KO seemed to require slightly less [Ca^2+^·Tcb2]_bound_ for growing fibers. D) Correlation of threshold for KOs to WT (% difference) vs time of delay (s). E) Model for EF-hands in Tcb2 self-assembly. C-terminal EF-hands is directly important for self-assembly while N-terminal EF-hands could be important for tuning/buffering Ca^2+^ for assembly or binding to other proteins.

Guided by this structural prediction, we used site-directed mutagenesis to introduce single alanine substitutions at each of the four putative calcium binding sites. We then expressed and purified each mutant (N-terminal: EF1mut (D37A), EF2mut (D79A); C-terminal: EF3mut (D141A), EF4mut (D184A)) to homogeneity. We then tested each mutant for qualitative network formation in our optical actuation assay using a strong test pulse (50μm, 10s, 100% intensity) (Fig. 5B, Movie S4). Mutations to the N-terminal EF-hand (EF1mut and EF2mut) were not disruptive to Tcb2 network formation and grew at similar rates and to similar sizes as WT Tcb2. In contrast, mutation to the C-terminal calcium binding site EF4mut showed no detectable network formation. Interestingly, although EF1-3 were competent for network formation, we noted qualitative differences in the microscopic appearance of the networks that formed. For example, EF2mut networks appeared more granular and disordered; while EF3mut displayed less contrast and appeared less granular. Together, these data show that the C-terminal EF-hand domain, and particularly the EF4mut binding site, are critical for calcium-dependent self-assembly of Tcb2 networks, while the N-terminal binding sites are non-essential, but potentially affect how monomers self-assemble.

To further characterize quantitative effects of the different N and C-terminal calcium binding sites on Tcb2 self-assembly, we used our combined Tcb2/Rhod-5N assay to monitor calcium loading into Tcb2 and network formation independently for each of the four EF hand mutants. For this purpose, we used a sensitive low-intensity long-duration pulse that leads to delayed network formation for WT Tcb2 (pulse parameters: 15s, 15%). As before, we tracked the maximum [Ca·Tcb2] in the field of view, and (where possible) radial network growth and the associated [Ca·Tcb2] at the network boundary (Fig. 5C). From this experiment we made several key observations. Based on the maximum [Ca·Tcb2], all four EF-hand mutants showed similar overall calcium loading dynamics during the pulse protocol, including the EF4mut that fails to form networks. This indicates that the other calcium binding sites in Tcb2-EF4mut are functional, further implicating calcium binding to EF4(D184) specifically as being critical to license self-assembly.

Second, although the other EF-hand mutants all supported network formation, they showed quantitative differences in the time-lag between pulse initiation and the appearance of Tcb2 networks. For the N-terminal EF hand domains, EF2mut formed networks more quickly than wildtype (WT 3.9s ± 0.9s vs. EF2mut 3.1s ± 0.6s, p=0.0261, n=10) and were able to grow further radially (WT 36.2μm ± 1.8μm vs EF2mut 41.7μm ± 3.28μm, p=0.0002, n=10); while, the EF1mut networks took slightly longer (Delay Time: WT 3.9s ± 0.9s vs. EF1mut 4.7s ± 0.5s, p=.0.0210, n=10) and grew slightly less radially (Max Growth Radius: WT 36.2μm ± 1.8μm vs. EF1mut 34.5μm ± 1.3μm, p=0.0318, n=10). In contrast, the C-terminal EF3mut took much longer to form networks (WT 3.9s±0.9s vs. EF3mut 6.1s±0.9s; p=0.0001, n=10) and grew to a much smaller network radius (WT 36.1μm ± 1.8μm vs. EF3mut 29.1μm ± 2.6μm; p=0.0001, n=10). These effects appeared to correlate with subtle differences in the calcium-loading requirements of each mutant at the network boundary (Fig. 5C, Fig. S3).

Collectively, the qualitative and quantitative measurements we performed on our panel of Tcb2 mutants define distinct roles for the N and C-terminal EF hand domains in controlling calcium-dependent Tcb2 self-assembly. The C-terminal EF hand appears to be the key module required for self-assembly: disruption of EF4mut completely abolished network formation, and disruption of EF3mut attenuated network formation significantly. In contrast, binding of calcium to the N-terminal EF hand is not critical for network formation. Indeed, elimination of the N-terminal EF2mut site actually increased Tcb2’s propensity to self-assemble. These results agree with single-domain NMR analyses which identified dramatic conformational changes upon binding of calcium ions to the C-terminal EF-hands specifically^34–36^ . Taken together, the C-terminal domain through the EF4mut site provides the key trigger for licensing Tcb2 self-assembly, while the N-terminal domains may act to tune the calcium responsiveness (Fig. 5D).

## Discussion

Here we have used a microscopy-based optical actuation assay to interrogate the biochemical requirements for calcium-regulated self-assembly of the cortical *Tetrahymena* cytoskeleton protein Tcb2. Using the photocaged calcium chelator DMNP-EDTA in combination with DMD patterned illumination, controllable and tunable calcium release dynamics could be generated and used to stimulate and monitor the self-assembly of Tcb2 networks in real time. By combining this approach with the low-affinity fluorescent calcium indicator Rhod-5N, we were able to infer the underlying spatiotemporal dynamics generated in this assay and estimate the associated loading of calcium into Tcb2 during assembly. This enabled us to identify a threshold concentration for calcium-bound Tcb2 needed to undergo a sharp ultrasensitive transition between soluble and self-assembled states. By applying this assay to Tcb2 mutants that disrupt distinct calcium binding sites in Tcb2’s N and C-terminal EF hand domains, we were able to identify EF4-D184 as the critical binding site that licenses Tcb2 for self-assembly, as well as suggest quantitative roles for other binding sites in tuning the protein’s calcium responsiveness.

Our approach and findings for the minimal one-component self-assembly of Tcb2 have implications for understanding other similar but more complex, multi-component calcium-regulated cytoskeletal systems. For example, many protists use structures called myonemes to achieve ultrafast calcium-regulated contractility—some capable of achieving extreme instantaneous cell velocities on the order of mm/s^19–23,26,45^. These myonemes typically contain centrin or spasmin-type calcium-binding proteins with similar EF-hand domain architectures to Tcb2. However, these proteins are typically physically arrayed onto so-called “giant proteins” such as Sfi1^17,18,21,23^, which serve as scaffolds containing hundreds of repeats of binding sites for individual centrin/spasmin monomers. Our optical actuation assay should allow for controlled interrogation of the calcium-responsiveness, spatiotemporal constraints, and contractility of these more complex structures, either through direct purification of myonemes from intact cells^22^ or through *in vitro* reconstitution of minimal systems.

Moreover, because some centrin-type homologues have also been observed to undergo Tcb2-like calcium-dependent self-assembly *in vitro*, one possible hypothesis for the contraction mechanism of myonemes is through constraining a Tcb2-like self-assembly process to occur along a pre-organized structure defined by the underlying scaffold protein^23,25,27^ . Our ability to use light-controlled calcium release to robustly control, probe, and tune the formation of Tcb2 networks from soluble components suggests an opportunity to test the plausibility of these biophysical mechanisms from the bottom-up using protein engineering and synthetic biology^30^ . Indeed, by creating synthetic scaffolding proteins that vary Tcb2 valency and spacing, it will likely be possible to quantitatively define and model the behavior of scaffolded self-assembly systematically. Such “synthetic myonemes” could potentially be harnessed for biotechnology applications to create customizable actuators with tunable mechanical properties and force generation capabilities.

While Tcb2 and related centrin-type proteins play structural and mechanical roles in the cell, their domain architecture and calcium binding activity make them similar to many proteins involved in cell signaling, such as calmodulin^2,44^. Like Tcb2, calcium binding to calmodulin licenses it to perform its downstream activities^1,3,46^. However instead of binding to itself, calcium-bound calmodulin typically interacts with and regulates the activity of a broad suite of other downstream targets, such as ion channels, motor-proteins, and kinases^2,46^. Interestingly, both experimental and theoretical studies have shown that calmodulin’s distinct N-terminal and C-terminal calcium-binding sites enable it to convert different patterns of spatiotemporal calcium activity—oscillations, transients, and bulk influx—into distinct signaling outcomes for cells^46–49^.

Our investigations into Tcb2 revealed a similar complexity as to how its EF-hands and calcium binding sites contribute to its interpretation of different spatiotemporal patterns for self-assembly. While the EF4 binding site was the most critical for licensing Tcb2 self-assembly, other binding sites appeared to alter the overall responsiveness which in turn affected the qualitative appearance of the networks that formed. Such effects may be critical to ensure Tcb2 structures assemble only under certain conditions or limit the growth and overall size of the resulting assemblies in living cells. Mechanistically, this may arise from buffering roles for certain sites, acting to sequester calcium away from the critical EF4 site and delay network formation. Alternatively, such binding sites might play roles in altering the conformational ensembles accessible to Tcb2, changing the nature of the self/self-interactions that support higher-order self-assembly. While a detailed understanding of the nanoscale interactions underlying Tcb2 assemblies awaits further investigation, our results have nonetheless revealed a surprisingly rich space of possible structures, responsiveness, and regulation available to a single-protein system mediated by its non-trivial interactions with calcium.

## Acknowledgements.

We thank members of the Coyle Lab, Bhamla Lab, A. Weeks, and M Elting for advice, helpful discussions, and critical reading of the manuscript. S.C. acknowledges support from NSF award 2313723 and the David and Lucille Packard Fellowship for Science and Engineering. J.H. acknowledges support from NSF award 2313727. S.B. acknowledges support from NSF award MCB-2313724 and National Institutes of Health (NIH) award R35GM142588.

## Methods

### Protein cloning, expression, purification

Full-length WildType Tcb2 DNA sequences were cloned into pET28 plasmid vectors with Kanamycin resistance. Primers were designed for single point mutations at each EF-hand domain. Plasmids were transformed into BL21 (DE3) E. Coli cells. Single colonies are inoculated into 50ml starter SOB cultures (with Kanamycin) overnight at 37°C shaking. Large cultures are inoculated with 10ml of starter cultures and are induced with 40mM IPTG once OD reaches 0.6-0.8 at 37°C then switched to 16°C after induction for overnight expression. Cells are harvested the next morning at 4000rcf for 20min and pellets are stored at -80°C for the next purification steps. Pellets are resuspended in B-PER Lysis Complete Reagent for lysis and sonicated for 2min 5on/off on ice and kept shaking in cold room for ∼30min. Lysate is clarified through centrifugation (10,000rcf, 20min). Pellets are washed 3 times with Inclusion Body Wash buffer (20mM Trish-HCl, 0.5M NaCl, 1mM EGTA, 1mM DTT, pH 7.2), sonicating for 1min 5on5off and clarifying at each wash step (10,000rcf, 20min). Pellets are washed one final time with MQ water, sonicated and clarified as before and pellets are stored in -80 till ready for next purification steps.

Wash frozen water pellets are washed in 25ml of Urea Wash buffer (4M Urea, 20mM MOPS, 100mM KCl, 1mM EGTA, 1mM DTT, pH 7.2) and clarified at 15,000xg for 20min. Urea wash supernatants are ultracentrifuged at 100,000xg for 1hr to remove and insoluble aggregates. Ultracentrifuged urea wash supernatants are dialyzed overnight (SnakeSkin Dialysis Tubing 3k MWCO), switched to fresh dialysis buffer, and dialyzed overnight again to remove urea (20mM MOPS, 100mM KCl, 1mM EGTA, 1mM DTT, pH 7.2). Dialyzed fractions are flash frozen as 1ml aliquots till ready for assays. For assays, 1ml aliquots of dialyzed fractions are thawed rapidly and are concentrated to a final concentration of ∼500μM using a centrifugal filter.

### Nikon Ti2-E Microscope Optical Configuration settings for optical actuation and calibration

All experiments are conducted using 40X objective with 405 LP filter on top turret and Multiband Filter in bottom turret. Solapad for wide-field fluorescence experiments is set to 5%. Auto-exposure is 40ms with 16bit camera sensitivity. The microscope is attached with a Mightex Polygon 1000-G DMD module for simultaneous multipoint photoactivation with different types of patterns, indicated here as regions of illumination (ROIs).

### DMNP-EDTA Caged calcium, Rhod-5N and Tcb2 stock preparation for optical actuation assays and calibration curves

DMNP-EDTA (Biotium, 5mg) is resuspended to a maximum concentration of 75mM in MQ Water and stored in -80. CaCl2 (75mM, MQ Water) is mixed with the 75mM DMNP-EDTA in a 1:1.3 ratio to make a 37.5mM caged calcium mix, then diluted to 1mM in MQ water. Rhod-5N (5mM, MQ Water) is diluted to 50µM (in dialysis buffer). 1mM caged calcium is mixed with 50µM Rhod-5N in 1:1 ratio (Final conc: 0.5mM caged calcium, 25µM Rhod-5N).

For optical actuation assays with Tcb2, 1μl of 5mM Rhod-5N is resuspended in 100ul of 500μM Tcb2 (final conc 50μM Rhod-5N). The Tcb2-Rhod stock is then similarly mixed in a 1:1 ratio with the 37.5mM caged calcium (Final conc: 0.5mM caged calcium, 250µM Tcb2, 25µM Rhod-5N).

### Dual Rhod-5N/Tcb2 optical actuation assay

1µl drops of the caged Ca-Rhod5N mix or caged Ca-Tcb2-Rhod mix are pipetted on to glass cover slides and the sample is prepared in a “sandwich structure” with a coverslip and a spacer (0.0035in thickness double sided tape) to maintain precise height for either brightfield or wide-field fluorescence microscopy. To initiate the release of Ca^2+^ from the DMNP-EDTA-Ca^2+^ complex, a 365 nm light pattern was projected through the Mightex Polygon 1000-G DMD. We define circular ROIs of different diameters (min: 10µm, max 200µm) and stimulate with varying pulse intensities (ranging from 10% to 100%) and pulse times (ranging from 1sec to 10sec). Time-lapse experiments are imaged at either 100ms or 200ms acquisition intervals for short stimulation profiles and 1sec or 2sec acquisition intervals for longer stimulation profiles. To correct for Rhod-5N bleaching downstream, we performed identical optical stimulation protocols on drops containing saturated Rhod-5N solution but without DMNP to obtain a timescale for intensity correction (Final conc: 50µM CaCl_2_, 25µM Rhod-5N).

### Rhod-5N calcium standard curves

Using Calcium Calibration buffer kit protocols as a guide, standards from 0.2mM - 50mM free Ca were made with 5uM Rhod-5N in each. 1ul drops of each standard were imaged using the same microscope acquisition settings as in the optical actuation assay. The average intensity of the center (1000×1000um region of analysis) of the drop is measured within ImageJ for each standard. Replicates of each standard were collected and plotted against the average intensities to build the calibration curve. Data points are fitted to the ‘two-parameter saturation growth” equation, y= (a*t)/(b+t); where parameter a is saturation level, parameter b is Kd for time, t.

### Image analysis and data processing

For Rhod-5N fluorescence images (in absence or presence of Tcb2), ND2 files loaded into Python and corrected for photobleaching. For the correction, we first generate a 2D spatiotemporal correction factor by dividing each frame of the saturated Rhod5N profiles by its first frame at a px level. The correction factor is then multiplied throughout the unsaturated Rhod5N profiles (in absence or presence of Tcb2). These raw intensity datasets are converted to local [Ca^2+^] by applying the standard curve equation to each 2D spatiotemporal profile for px level quantification.

For Tcb2 brightfield images, a contrast-based pipeline is used to detect and quantify the radial growth of self-assembling Tcb2 networks. Time-lapse experiments (.nd2 file format) are loaded and analyzed as stacks using FiJI/ImageJ. Using the “Find Edges” plugin, stacks are converted to rate of contrast intensity change by a Sobel Edge detector. A Gaussian Blur at 5px radius is applied and a threshold is set to create a binary mask of the fiber assembly. Within Python, we find the largest points away from the center for each frame and fit each frame to a circle to get the radial growth of the fiber assembly.

## Notes

### Competing Interest Statement

The authors have declared no competing interest.

