## Supplemental Figures for "Decoding ultrasensitive self-assembly of the calcium-regulated *Tetrahymena* cytoskeletal protein Tcb2 using optical actuation"

**Affiliations:**
[1]Department of Biochemistry, University of Wisconsin-Madison
[2]Integrated Program in Biochemistry Graduate Program
[3]School of Chemical and Biomolecular Engineering, Georgia Institute of Technology
[4]Department of Biology, Drake University

University of Wisconsin-Madison, Madison, Wisconsin 53706, USA.

Contains:

Supplemental figures 1-3.

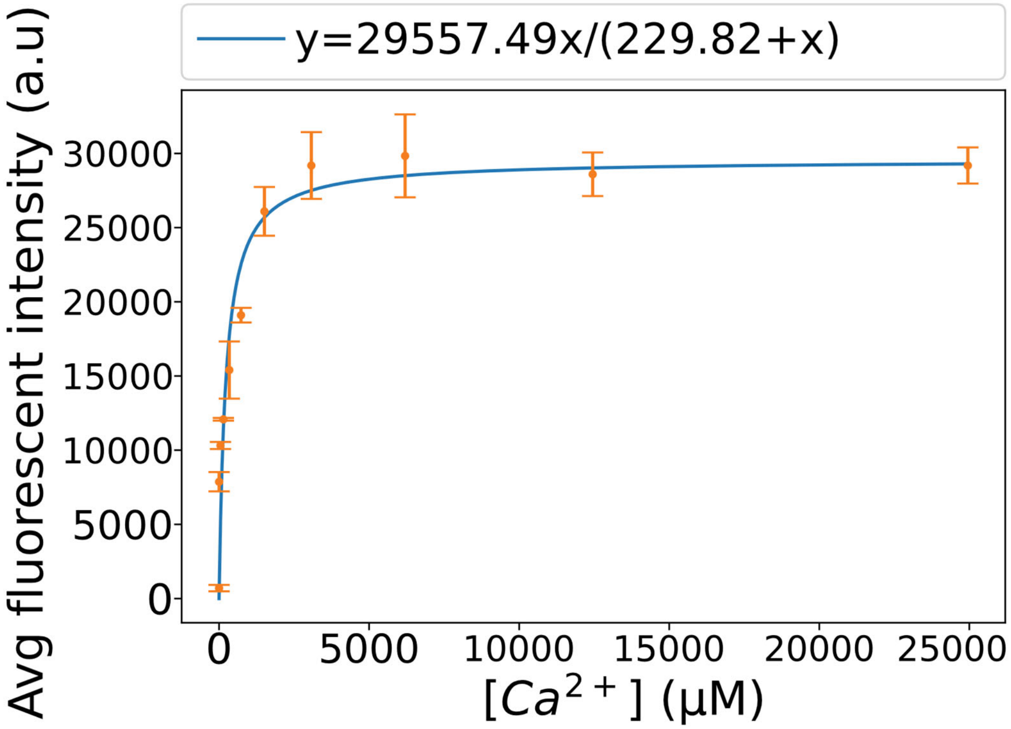


**Supplemental Figure 1. Sample calibration curve equation determine fitting standards of 2µM to 25mM [$Ca^{2+}$] in the presence of 25uM Rhod-5N imaged with our optical set up.** Standards are fitted to a binding curve (y=A*x/$K_D$ + x) and rearranged to make a calibration curve that converts Rhod-5N intensities into calcium concentrations.

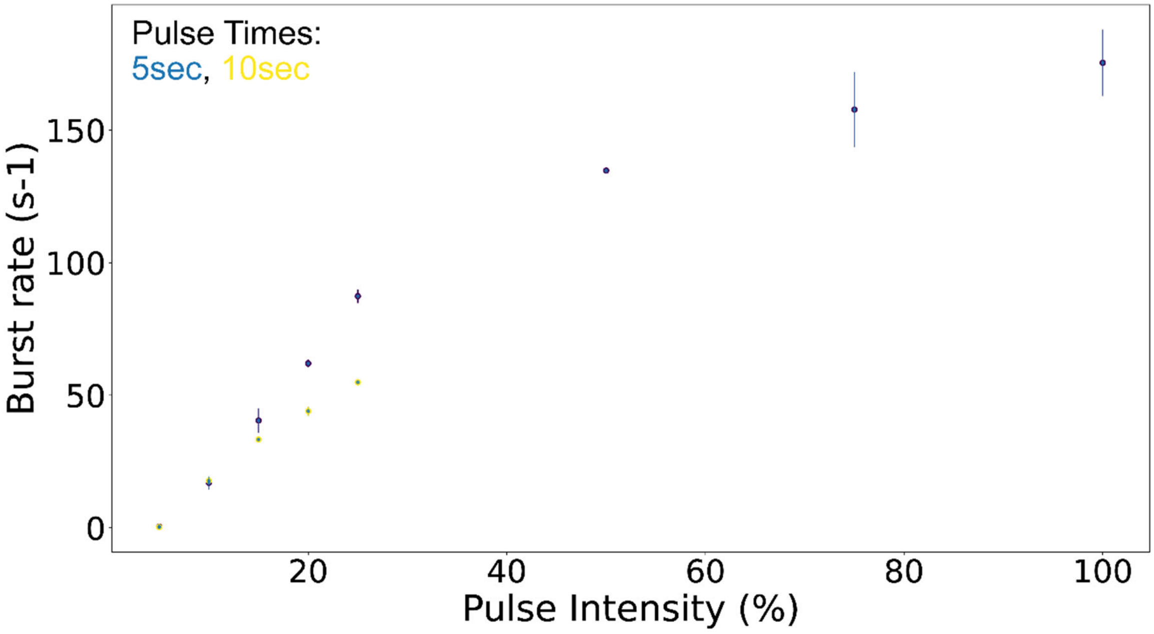


**Supplemental Figure 2: Rate of burst phases scales with intensity**

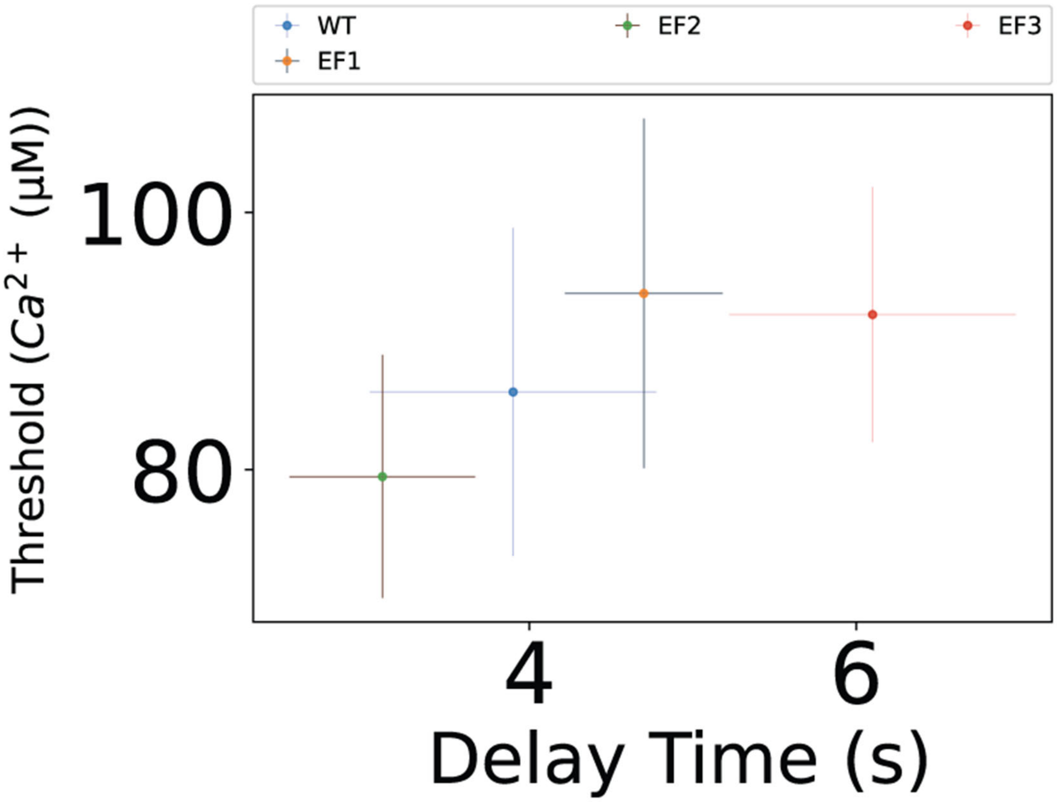


**Supplemental Figure 3. Correlation between [$Ca^{2+}$·Tcb2] threshold and time delay in triggering Tcb2 self -assembly among the EF-hand knockouts.** Lowered threshold correlates with a faster time delay while a higher threshold indicates a longer time delay.
